# Genetic tagging uncovers a robust, selective activation of the thalamic paraventricular nucleus by adverse experiences early in life

**DOI:** 10.1101/2022.12.05.519218

**Authors:** Cassandra L. Kooiker, Yuncai Chen, Matthew T. Birnie, Tallie Z. Baram

**Affiliations:** Department of Anatomy and Neurobiology, University of California- Irvine, Irvine, CA; Department of Pediatrics, University of California- Irvine, Irvine, CA; Department of Neurology, University of California- Irvine, Irvine, CA

**Author notes:** Corresponding author: Tallie Z. Baram, MD, PhD, Medical Sciences B, ZOT 4475; UCI, Irvine CA 92697-4475.

**Keywords:** genetic tagging, TRAP2, PVT, reward circuit, stress, ELA, Fos

## Abstract

**Background:** Early-life adversity (ELA) is associated with increased risk for mood disorders including depression and substance use disorders. These are characterized by impaired reward-related behaviors, suggesting compromised operations of reward-related brain circuits. However, the brain regions engaged by ELA that mediate these enduring consequences of ELA remain largely unknown. In an animal model of ELA, we have identified aberrant reward-seeking behaviors, a discovery that provides a framework for assessing the underlying circuits.

**Methods:** Employing TRAP2 male and female mice, in which neurons activated within a defined timeframe are permanently tagged, we compared ELA and control-reared mice, assessing the quantity and distribution of ELA-related neuronal activation. After validating the TRAP2 results using native cFos labeling, we defined the molecular identity of this population of activated neurons.

**Results:** We uniquely demonstrate that the TRAP2 system is feasible and efficacious in neonatal mice. Surprisingly, the paraventricular nucleus of the thalamus (PVT) is robustly and almost exclusively activated by ELA and is the only region distinguishing ELA from typical rearing. Remarkably, a large proportion of ELA-activated PVT neurons express CRFR1, the receptor for the stress-related peptide, corticotropin-releasing hormone (CRH), but these neurons do not express CRH itself.

**Conclusions:** We show here that the PVT, an important component of reward circuits which is known to encode remote, emotionally salient experiences to influence future motivated behaviors, encodes adverse experiences as remote as those occurring during the early postnatal period and is thus poised to contribute to the enduring deficits in reward-related behaviors consequent to ELA.

## Introduction

Early life adversity consisting of trauma, poverty, or tumultuous environment impacts the lives of over 30% of children in the United States (1). In humans, ELA is associated with poor cognitive and emotional health and increased risk for mood disorders such as depression as well as increased risk for substance use disorders (2-6). Human imaging studies suggest altered development of specific brain circuits following ELA, including reward circuits (7,8). It is crucial to understand the nature of these associations because ELA and its consequences, unlike genetic contributors to vulnerability to psychopathologies, may be amenable to prevention. In human studies, it is difficult to demonstrate causality between ELA and adverse adult outcomes and to establish the underlying mechanisms, necessitating use of experimental animal models. Using an animal model in which ELA is induced by an impoverished environment that provokes aberrant maternal care, we have identified later life disruptions of reward-related behaviors (9-13). These disruptions indicate dysfunction of the reward circuit, as seen in several human mood disorders that commonly follow ELA (5,14). These findings provide an impetus to employ this animal model for advancing our understanding of how transient ELA enduringly disrupts the operations of reward circuits.

Here we aimed to determine which regions and neuronal populations within the brain, and specifically within reward circuits, were activated by ELA and are thus candidates for mediating the behavioral consequences of ELA. To this end, we employed TRAP2 mice expressing iCre-ER^T2^ recombinase at the locus of the immediate early gene, cFos, to genetically label neurons that are activated during ELA with the red fluorescent reporter tdTomato using Ai14, a knock-in allele of the Rosa26 locus (15). To date this technique had not been utilized during the first week of life. Using this approach, we established the paraventricular nucleus of the thalamus (PVT) as a key region engaged by these experiences.

The PVT is a dorsal midline thalamic nucleus that is a crucial component of the limbic system and emotional processing network and is engaged by emotionally salient stimuli of either valence (16, 17). The PVT utilizes information derived from remote emotionally salient experiences to gate the expression of approach and avoidance behaviors and to influence responses to stress (17-21). The PVT projects to many brain regions with important roles in stress and reward (22-24) and is thus poised to regulate behaviors related to stress and reward following salient early life experiences. Indeed, our prior work using cFos expression had indicated that the PVT is activated by positive early life experience in the form of augmented maternal care (25), yet, whether this applies to experiences of the opposite valence and whether this activation may influence future behaviors remains unknown.

## Methods and Materials

### Animals

Fos^2A-iCreER^ (Jax #030323) and Ai14 (Jax #007914) mice were received from The Jackson Laboratory or bred in house. All mice were housed in a temperature-controlled, quiet and uncrowded facility on a 12-hr. light, 12 hr. dark schedule (lights on at 0630 hr., lights off at 1830 hr.), except Fos^2A-iCreER^+/+ litters. These litters were maintained on a 12-hr. reverse light cycle (lights on at 0000 hr., lights off at 1200 hr.) to enable perfusion during the mice’s early active period, and we have previously found no difference in any behavioral test between normally-housed and reverse-light-cycle-housed mice when tested at the same point of their circadian cycle. Mice were provided with *ad libitum* access to water and food (Envigo Teklad, 2020x, global soy protein-free extruded). Fos^2A-iCreER^ mice bred with the Ai14 reporter mice were employed for TRAP2 studies; Fos^2A-iCreER^+/+ were employed for the endogenous cFos validation studies. All experiments were performed in accordance with National Institutes of Health guidelines and were approved by the University of California-Irvine Animal Care and Use Committee.

### The Limited Bedding and Nesting (LBN) Model of Early-life Adversity (ELA)

Fos^2A-iCreER^ dams bred with Ai14 males were singly housed on embryonic day (E)17 and monitored for birth of pups every 12 hours. On the morning of postnatal day (P)2, Fos^2A-iCreER^+/-::Ai14+/- litters and Fos^2A-iCreER^+/+ litters were culled to a maximum of 8 pups, including both sexes, and the ELA paradigm was initiated as previously described (26). Control dams and pups were placed in cages with a standard amount of corn cob bedding (400 ml) and cotton nestlet material (one 5×5 cm square). ELA dams and pups were provided with one-half cotton nestlet placed on a 2.5 cm tall, fine gauge plastic-coated aluminum mesh platform above sparse corn cob bedding on the cage floor. Accumulation of ammonia was avoided by placing cages in a room with robust ventilation. Both rearing groups were left completely undisturbed until the morning of P6, at which point pups were briefly removed from the cage, placed on a warming pad, and injected subcutaneously with 150 mg/kg tamoxifen (cat#T5648, Millipore Sigma) dissolved in corn oil (cat#C8267, Millipore Sigma; Fig. S1). Pups were then returned to their cage and left undisturbed until the morning of P10. Dams and pups were then transferred to standard cages where maternal behaviors rapidly normalize and the ELA pups’ stress dissipates. P6 was chosen as the timepoint for tamoxifen injections because it is roughly at the midpoint of the ELA period and should allow for optimal tagging of neurons activated by ELA rearing.

### Brain processing and analyses

On the morning of P14 (for Fos^2A-iCreER^+/-::Ai14+/- litters), or at approximately 1400 hr. of P10 (for Fos^2A-iCreER^+/+ litters), the dam was removed from the home cage, and the cage was placed on a warming pad. Pups were euthanized with sodium pentobarbital and transcardially perfused with ice cold phosphate-buffered saline (PBS; pH=7.4) followed by 4% paraformaldehyde in 0.1M sodium phosphate buffer (pH=7.4). Perfused brains were post-fixed in 4% paraformaldehyde in 0.1M PBS (pH=7.4) for 4-6 hrs. before cryoprotection in 25% sucrose in PBS. Brains were then frozen and coronally sectioned at a thickness of 30 μm (1:5 series) using a Leica CM1900 cryostat (Leica Microsystems, Wetzlar, Germany). Fos^2A-iCreER^+/-::Ai14+/- sections were mounted on gelatin-coated slides and coverslipped with Vectashield containing DAPI (cat. #H-1200, Vector Laboratories, Burlingame, CA, USA). P14 was chosen as the sacrifice timepoint for Fos^2A-iCreER^+/-::Ai14+/- litters because optimal expression/accumulation of the tdTomato reporter is not achieved until at least a week following the tamoxifen injection.

### Immunohistochemistry

Avidin-biotin complex (ABC)-amplified, diaminobenzidine (DAB) reactions were used to visualize cFos and CRFR1 on free-floating sections. Sections were first washed in PBS containing 0.3% triton (PBST; 3 × 5 min.) followed by quenching of endogenous peroxidase activity by incubation in 0.3% H_2_O_2_ for 20 min. Sections were blocked in 5% normal donkey serum (NDS) or normal goat serum (NGS) in PBST for one hr. Sections were incubated with 1:40,000 rabbit anti-cFos (cat# ABE457, Millipore Sigma, Temecula, CA) for 3 days at 4°C or 1:2,000 goat anti-CRFR1 (cat# EB08035, Everest Biotech, Ramona, CA) for 16 hrs. at 4°C. Following 3 × 5 min. washes in PBST, sections were incubated with 1:500 biotinylated goat anti-rabbit antibody (cat# BA-1000-1.5, Vector Laboratories, Burlingame, CA) or 1:500 biotinylated donkey anti-goat antibody (cat# 705-065-147, Jackson ImmunoResearch, West Grove, PA) in 2% NDS or NGS for 2 hrs. Sections were washed in PBST (3 × 5 min.) and then incubated in 1% ABC solution (Vectastain, Vector Laboratories, Burlingame, CA) and washed again in PBST (3 × 5 min.). The immunoreaction product was visualized using solution containing 0.005% H_2_O_2_ and 0.05% DAB. Sections were mounted onto gelatin-coated slides or co-labeled for tdTomato expression.

For co-labeling to visualize the cFos reporter, tdTomato, benzidine dihydrochloride (BDHC) reactions were used following DAB staining. Sections were quenched in a solution of 50% methanol and 0.2% H_2_O_2_ in PBST for 5 min. followed by 100% methanol containing 0.2% H_2_O_2_ for 20 min. Sections were washed in PBST (3 × 5 min.) then blocked in 2% NGS for 30 min. Sections were then incubated with 1:10,000 rabbit anti-RFP (cat# 600-401-379, Rockland Immunochemicals, Pottstown, PA) for 3 days at 4°C. Following primary antibody incubation, sections were washed in PBST (3 × 5 min.) and incubated in 1:500 biotinylated goat anti-rabbit antibody in 2% NGS for 2 hrs. Sections were then washed in PBST (3 × 5 min.) and incubated in 1% ABC solution, followed by additional washes in PBST (3 × 5 min.). Sections were washed in 1X acidic buffer (3 × 5 min.; cat# 003850, Bioenno, Santa Ana, CA) then incubated in a buffer containing 0.025% sodium nitroprusside and 0.01–0.02% benzidine dihydrochloride (BDHC) for 5–10 min. The granular blue deposits were visualized by immersing the sections in fresh incubation solution containing 0.003% H_2_O_2_ for 3 min. Sections were washed in 1X acidic buffer (3 × 5 min.) and mounted onto gelatin-coated slides. All mounted sections were dehydrated and coverslipped with Permount mounting medium (cat# SP15-500, Fisher Scientific, Hampton, NH).

For fluorescent labeling of CRH, the tyramide signal amplification technique was used (27). Free-floating sections were blocked in 5% NGS in PBST for one hour and then incubated in rabbit anti-CRH antiserum (1:20,000; gifted by Dr. W. Vale, Salk Institute, La Jolla, CA) for 14 days at 4°C. Following washing in PBST (3 × 5 min.), sections were incubated in horseradish peroxidase-conjugated anti-rabbit IgG (1:1000; cat# NEF812001EA, Perkin Elmer, Boston, MA) for 1.5 hours. Fluorescein tyramide conjugate was diluted in in amplification buffer (1:150; cat# NEL701A001KT, Perkin Elmer, Boston, MA) and applied to sections in the dark for 5-6 min., followed by washing and mounting on gelatin-coated slides. Sections were coverslipped with Vectashield containing DAPI.

### Image acquisition

Images of sections processed with DAB and BDHC were collected using a Nikon Eclipse E400 light microscope with 10x and 20x objective lenses. Confocal images were collected using an LSM-510 confocal microscope (Zeiss, Dublin, CA, USA) with an Apochromat 10x, 20x, or 63x objective. Virtual z-sections of 1 μm were taken at 0.2–0.5 μm intervals. Image frame was digitized at 12-bit using a 1024 × 1024-pixel frame size.

### Analyses and statistical considerations

tdTomato^+^, CRFR1^+^, CRH^+^, and cFos^+^ neuron numbers were counted manually in FIJI (28). All quantifications and analyses were performed using stereological principles including systematic unbiased sampling and without knowledge of group assignment. Statistical analyses were carried out using GraphPad Prism software (GraphPad, San Diego, CA, USA). Differences between CTL and ELA groups of both sexes were assessed using two-way ANOVA. To examine significance of cell number differences throughout the anteroposterior axis of the PVT, we used two-way mixed ANOVA with the Sidak multiple comparisons *post hoc* test.

## Results

### TRAP2 system is feasible and effective in neonatal mice

We utilized TRAP2;Ai14 mice expressing iCre-ER^T2^ recombinase in an activity-dependent manner to initiate expression of an iCre-dependent tdTomato (tdT) reporter. To initiate recombination and reporter expression, tamoxifen was administered to postnatal day 6 (P6) pups reared in standard or ELA cages, and reporter expression was assessed one week later to allow for optimal accumulation of the reporter. Modest, consistent tdT expression was detected in cell bodies in several brain regions in ELA mice, including the hypothalamic paraventricular (PVN) and suprachiasmatic nuclei (SCN; Fig. 1). The most robust and striking reporter expression, indicative of neuronal activation during the P6-P8 time period, was identified in the paraventricular nucleus of the thalamus (PVT).

**Fig. 1.**
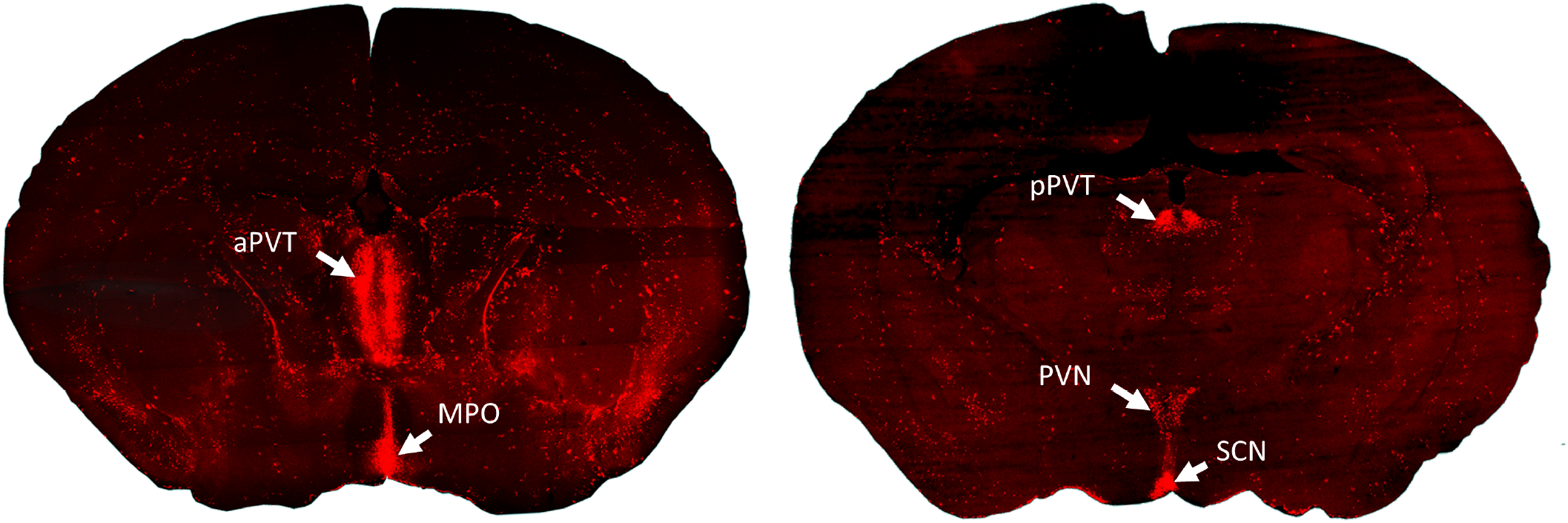
PVT neurons are robustly and selectively activated during exposure to adverse rearing conditions. Low magnification images of brain sections from TRAP2 mice sacrificed a week after receiving tamoxifen on P6 to enable cFos-dependent Cre expression for 24-48 hrs. Strong activation of the PVT, with little reporter expression in the rest of the brain is apparent. aPVT, anterior paraventricular nucleus of the thalamus; MPO, medial preoptic nucleus; pPVT, posterior paraventricular nucleus of the thalamus; PVN, paraventricular nucleus of the hypothalamus; SCN, suprachiasmatic nucleus.

### PVT neurons are selectively activated by early life adversity

It has been established that, in adults, the PVT is activated by emotionally salient stimuli of both positive and negative valence (16,17). Here, we sought to determine if PVT activation occurred during emotionally salient experiences during the neonatal/infancy period. To this end, we reared mice either in typical cages or in our model of early life adversity (ELA). In this simulated poverty model, mouse pups are reared in cages with limited bedding and nesting materials, conditions which provoke aberrant maternal care behaviors and stress in the pups (26,29-30). Exposure to this environment from P2 to P9 induces persistent disruptions in reward-related behaviors later in life (12,13). We administered tamoxifen to both control and ELA pups on P6, which results in induction of reporter expression in all neurons activated during a 24–48-hour time window following tamoxifen administration.

There was very little neuronal activation, measured by number of neurons expressing tdT, throughout the brains of P6-8 male and female mice (Figure 1). However, a screen of serial coronal sections throughout the brain suggested that prominent cFos expression took place in the PVT in both control and ELA mice. Analyzing male mice, a two-way ANOVA with Sidak’s *post hoc* test identified significantly greater overall PVT activation in ELA males as compared to control males (p = 0.002, Fig. 2A,B,E). A two-way mixed model ANOVA with coordinate as a repeated factor identified a main effect of rearing on reporter expression (p<0.001), and Sidak’s multiple comparison’s *post hoc* test indicated a significantly larger number of TRAPed (tdT^+^) neurons in ELA males as compared to control males at several coordinates along the anteroposterior axis of the PVT (at -0.46 mm, -1.22 mm, -1.46 mm, and -1.7 mm from Bregma), indicating that this differential activation was particularly prominent in the mid to posterior PVT (Fig. 2H). Comparison of overall PVT activation between control males and control females demonstrated a greater number of TRAPed PVT neurons in control females (p = 0.019), and this was particularly prominent in the posterior PVT (−1.94 mm from Bregma; p = 0.007; Two-way mixed model ANOVA with Sidak *post hoc* test; Fig. 2E,F). Strikingly, and in contrast to males, ELA did not further augment the density of TRAPed cells in the PVT of female mice (Fig. 2C, D, I).

**Fig 2.**
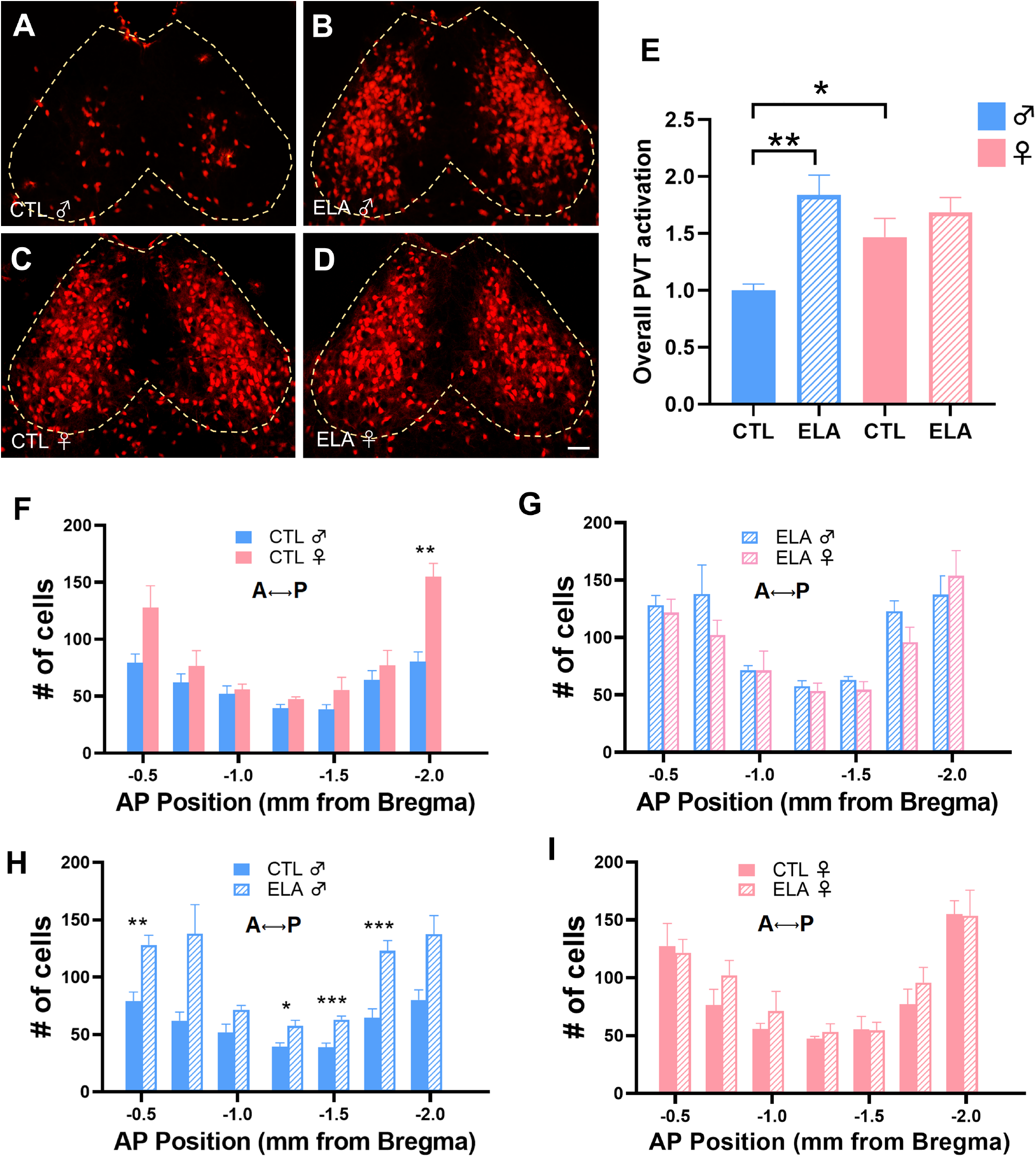
ELA induces greater PVT activation in males compared with females. (**A**) Control and (**B**) early life adversity (ELA) P14 male TRAP2 mouse reporter (tdT) expression in the pPVT following tamoxifen administration on P6. (**C**) Control and (**D**) ELA P14 female TRAP2 mouse tdT expression in the pPVT following tamoxifen administration on P6. Scale bar = 50 μm. (**E**) Comparison of overall PVT activation in male and female control and ELA mice, normalized to activation in control males (n = 7-9 mice/group; Two-way mixed model ANOVA with the Sidak *post hoc* test). (**F**) Quantification of tdT^+^ neurons across the anteroposterior axis of the PVT comparing control males and females. Two-way mixed model ANOVA with the Sidak *post hoc* test shows significantly more TRAPed neurons at -1.94 mm from Bregma in females (p = 0.007; n = 7-10 mice/group). (**G**) Comparing the number of tdT^+^ PVT neurons in ELA males and females indicates no difference (n = 8-14 mice/group; (p = 0.296; Two-way mixed model ANOVA). (**H**) Two-way mixed model ANOVA with the Sidak *post hoc* test comparing control and ELA males indicates significantly more tdT^+^ PVT neurons at -0.46, -1.22, -1.46, and -1.70 mm from Bregma (n = 10-14 mice/group; Two-way mixed model ANOVA with the Sidak *post hoc* test). (**I**) Comparing the number of tdT^+^ PVT neurons in control and ELA females indicates no difference (p = 0.472; n = 7-8 mice/group; Two-way mixed model ANOVA). ***, p < 0.001; **, p < 0.01; *, p < 0.05. All values shown as mean ± SEM.

Reporter expression in TRAP2 mice is driven by the cFos promoter and aims to reflect cFos expression as a marker of neuronal activation. Because the use of the TRAP2 transgenic system during the first week of life has not yet been published, we determined the validity of this method during the neonatal period by visualizing native cFos expression in the PVT during control and ELA rearing conditions. Immunolabeling against cFos in the PVT of P10 mice was congruent with that of the TRAP2 reporter (Fig. 3). As expected, significantly more cFos^+^ neurons were found in the PVT of ELA males as compared to control males. In accord with the TRAP2 reporter expression, we found no difference in number of cFos^+^ neurons when comparing control and ELA females. These data support the conclusion that the use of the TRAP2 system in early life produces reporter expression reflective of native cFos expression during this period.

**Fig 3.**
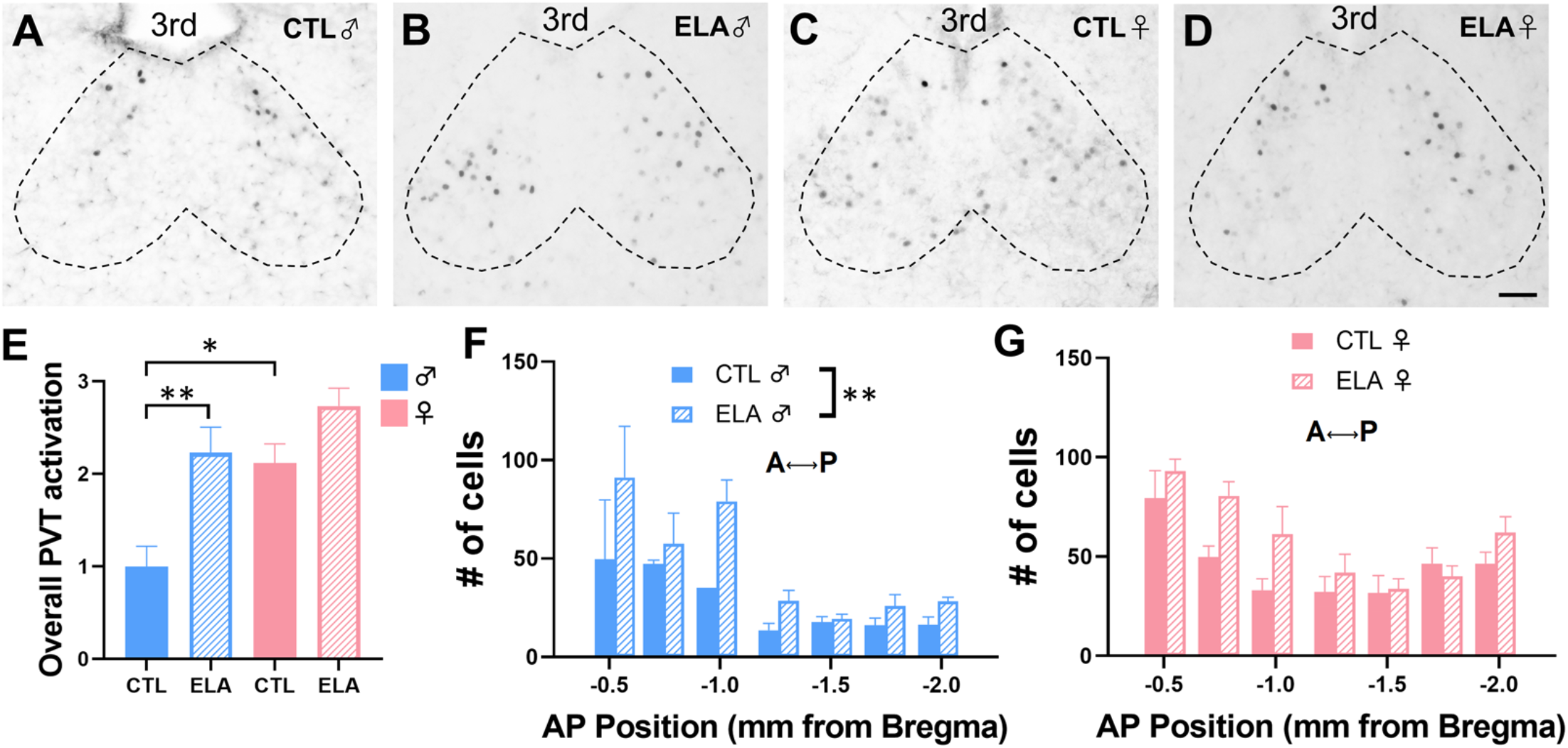
Endogenous cFos expression in the PVT is congruent with and validates the TRAP2 method. (**A**) Representative images of cFos expression in the pPVT of P10 control males, (**B**) ELA males, (**C**) control females, (**D**) and ELA females sacrificed immediately following exposure to normal or adverse rearing conditions; 3^rd^, third ventricle. Scale bar = 50 μm. (**E**) Comparison of overall PVT cFos expression in male and female control and ELA mice, normalized to activation in control males (n = 5-6 mice/group; Two-way mixed model ANOVA with the Sidak *post hoc* test). (**F**) Quantification of cFos^+^ neurons across the anteroposterior axis of the PVT in P10 control and ELA male mice (**G**) and female mice (n = 7-11 mice/group; Two-way mixed model ANOVA). **, p < 0.01; *, p < 0.05. All values shown as mean ± SEM.

### Increased neuron activation by early life adversity in males is selective to the PVT

Quantification of TRAPed neurons in control and ELA males was performed in several additional brain regions, including those involved in reward and responses to stress, such as the nucleus accumbens, amygdala, ventral tegmental area, and paraventricular nucleus of the hypothalamus (PVN). This quantification largely revealed sparse reporter expression in both rearing groups, suggesting little activation during P6-P8 in most regions outside of the PVT (Fig. 4, Table 1). Importantly, in all areas analyzed besides the PVT, reporter expression did not differ significantly between ELA and control male mice (Table 1). These findings indicate that the augmented number of TRAPed neurons in the PVT of ELA males as compared to control males is specific to this nucleus.

**Fig 4.**
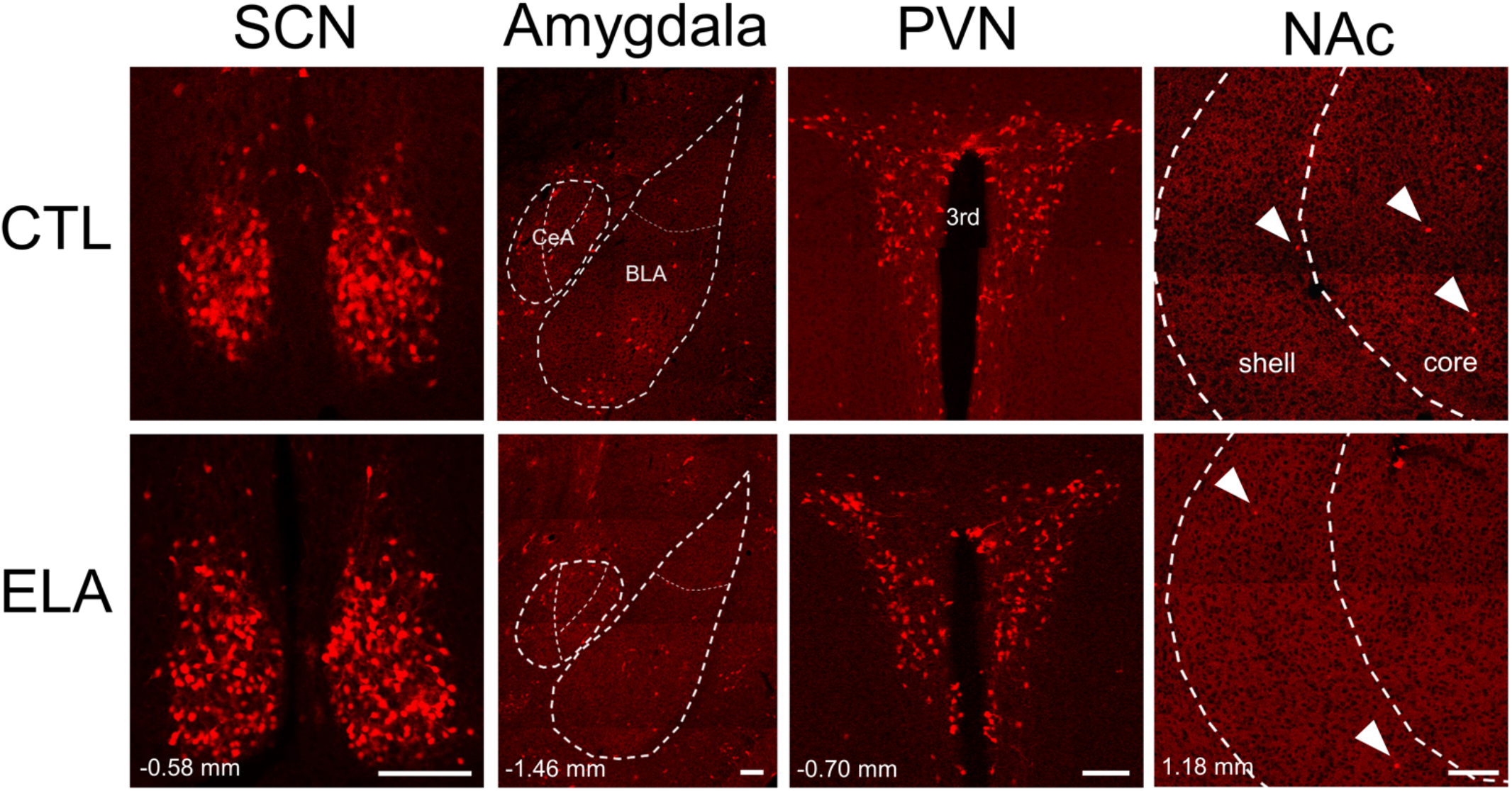
The numbers of TRAPed neurons differ between control and ELA male mice exclusively in the PVT. Representative images for multiple brain regions show no difference in tdTomato expression in male TRAP2 mice reared in control vs. ELA conditions. Few tdTomato^+^ neurons are found in most regions, including those important in reward- and stress-related behaviors. Coordinates indicate the distance from Bregma of each image. SCN, suprachiasmatic nucleus; CeA, central nucleus of the amygdala; BLA, basolateral amygdala; PVN, paraventricular nucleus of the thalamus; 3^rd^, third ventricle; NAc, nucleus accumbens. Scale bars = 100 μm.

**Table 1.** Region-specific quantification of tdTomato^+^ neurons in male P14 TRAP2 mice TRAPed during normal rearing or adverse rearing conditions, with 95% confidence intervals.

| Brain region | CTL TRAPed<br>Neurons | ELA TRAPed<br>Neurons |  |
| --- | --- | --- | --- |
| Paraventricular thalamus<br>Anterior<br>Mid<br>Posterior | 79.31 ± 18.57<br>39.33 ± 7.94<br>80.31 ± 20.75 | 133.60 ± 16.80<br>57.65 ± 10.92<br>145.10 ± 35.40 | p = 0.0001<br>p = 0.008<br>p = 0.005 |
| Nucleus accumbens<br>Core<br>Medial shell | 9.16 ± 2.49<br>1.73 ± 0.65 | 7.76 ± 2.58<br>1.63 ± 0.73 | p = 0.397<br>p = 0.823 |
| Amygdala<br>Basolateral<br>Central | 17.00 ± 3.99<br>11.29 ± 6.18 | 13.61 ± 5.24<br>8.90 ± 6.31 | p = 0.278<br>p = 0.556 |
| Medial prefrontal cortex<br>Prelimbic<br>Infralimbic | 5.29 ± 1.03<br>7.13 ± 3.64 | 6.88 ± 5.24<br>6.14 ± 2.95 | p = 0.521<br>p = 0.631 |
| Ventral tegmental area | 4.80 ± 2.54 | 6.11 ± 3.85 | p = 0.516 |
| Hypothalamus<br>Suprachiasmatic<br>Paraventricular<br>Lateral<br>Ventromedial | 56.00 ± 9.44<br>21.67 ± 7.92<br>12.14 ± 7.03<br>6.71 ± 4.06 | 65.57 ± 10.81<br>25.58 ± 9.34<br>10.43 ± 5.47<br>3.43 ± 2.61 | p = 0.137<br>p = 0.489<br>p = 0.646<br>p = 0.121 |
| Hippocampus<br>CA1<br>CA3<br>Dentate gyrus | 14.33 ± 5.69<br>10.17 ± 3.48<br>20.50 ± 9.97 | 11.75 ± 2.95<br>10.38 ± 3.82<br>19.13 ± 6.93 | p = 0.301<br>p = 0.926<br>p = 0.778 |

### Molecular/phenotypic characterization of ELA-TRAPed PVT neurons

The PVT is a heterogenous structure composed of numerous cell types that are defined by the expression of distinct neurotransmitters, neuropeptides, and receptors. Hence, we next sought to determine the molecular characteristics of PVT neurons engaged by early-life experiences, including ELA, to gain insight into the potential functional roles of this population. In addition, because ELA intrinsically involves stress, and neurons expressing the stress-related neuropeptide CRH are impacted by ELA in the hypothalamus (31,32), amygdala (10,33), and hippocampus (34), we focused on PVT neurons expressing CRH or its receptors. Immunostaining against CRH demonstrated a rich population of CRH-expressing neurons in the PVT, in accord with prior reports (35,36).

However, no TRAPed PVT neurons expressed CRH (Fig. 5A, B). In contrast, immunostaining against the type 1 receptor to CRH, CRFR1, demonstrated a major increase in the proportion of activated PVT neurons expressing CRFR1 in ELA mice as compared to controls. Specifically, in males, 40.8% of TRAPed PVT neurons expressed this receptor in ELA mice (95% CI [0.349. 0.468]), whereas 20.3% of TRAPed PVT neurons expressed this receptor in control males (95% CI [0.151, 0.255]). In females, 50.5% of TRAPed PVT neurons expressed CRFR1 in ELA mice (95% CI [0.441, 0.569]), and 24.5% of TRAPed PVT neurons expressed CRFR1 in controls (95% CI [0.185, 0.304]). This effect was particularly prominent in the anterior and mid PVT in females and across the entire anteroposterior axis of the PVT in males (Fig. 5E, F). Therefore, in both males and females, a significantly larger proportion of neurons engaged in early life expressed CRFR1 in ELA groups as compared to controls. Additionally, two-way mixed model ANOVA comparing the proportion of TRAPed aPVT neurons that express CRFR1 in control and ELA males and females indicated a significant interaction between rearing condition and sex (p = 0.002; Fig. S2), with females experiencing a larger ELA-induced increase in proportion of TRAPed aPVT neurons expressing this receptor. Surveys of CRFR2 expression in PVT using both *in situ* hybridization (37) and IHC revealed only low levels, so this receptor was not quantified.

**Fig 5.**
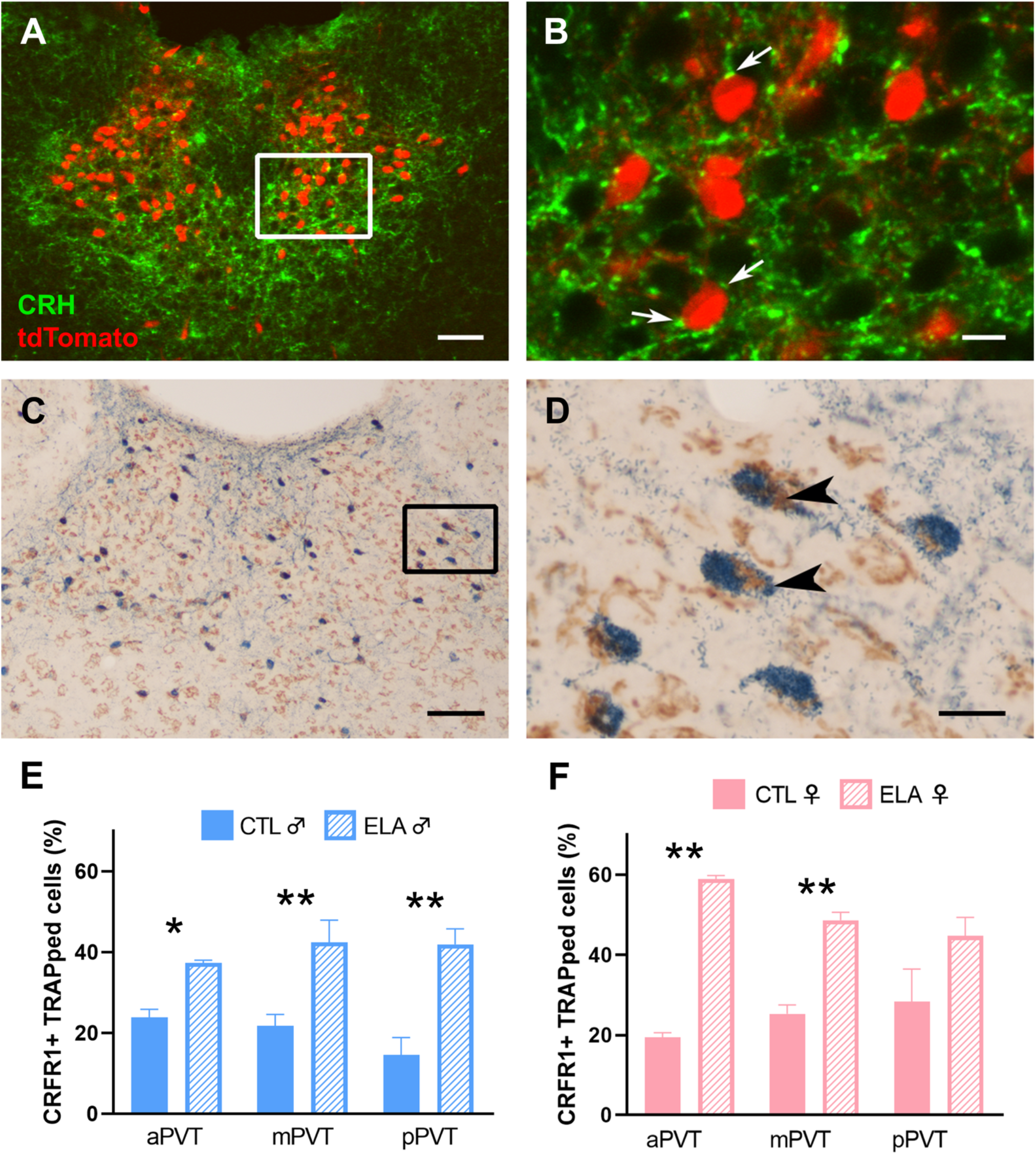
TRAPed PVT neurons express CRFR1 but not CRH. (**A**) Representative image from the pPVT of tdTomato (red) and CRH (green) expression in a P14 TRAP2 male demonstrating lack of overlap between CRH and the TRAP2 reporter. Scale bar = 50 μm. (**B**) Higher magnification reveals numerous CRH^+^ puncta around TRAPed PVT neurons (white arrows) but no CRH expression within TRAPed neurons. Scale bar = 10 μm. (**C**) Representative image from the pPVT of tdTomato (blue) and CRFR1 (brown) expression in a P14 TRAP2 mouse, showing robust overlap of CRFR1 and reporter expression. Scale bar = 50 μm. (**D**) Higher magnification reveals partial co-expression of tdTomato and CRFR1. Scale bar = 10 μm. (**E**) In males, and across anterior, mid, and posterior PVT, a significantly larger proportion of TRAPed PVT neurons express CRFR1 following ELA as compared to control rearing (n = 6 mice/group; Two-way mixed model ANOVA with the Sidak *post hoc* test). (**F**) In females, across anterior and mid PVT, a significantly larger proportion of TRAPed PVT neurons express CRFR1 following ELA as compared to control rearing (n = 5-6 mice/group; Two-way mixed model ANOVA with the Sidak *post hoc* test). **, p < 0.01; *, p < 0.05. All values shown as mean ± SEM.

## Discussion

The principal findings in this set of experiments are (1) genetic tagging of neurons activated during the neonatal period in mice is feasible, with high sensitivity and fidelity; (2) the PVT is the major brain region activated by early-life adversity; (3) sex is an important determinant of neuronal activation by early-life experiences, and (4) neurons expressing the CRH receptor, CRFR1, likely a target of CRH signaling, are preferentially activated by ELA in a sex-dependent manner and are poised to contribute to the mechanisms by which it contributes to alterations in adult behaviors.

Using the TRAP2 transgenic mouse, we identify here region-specific neuronal activation during the early postnatal period in the mouse. Reporter expression is highly congruent with native cFos. Unlike native cFos expression, the TRAP2 system allows for labeling of neuronal activity over a much longer time period (up to approximately 48 hours) following tamoxifen administration (15). This characteristic is advantageous when visualizing neuronal activity during a chronic stimulus, such as early-life adversity. The finding that activity-dependent genetic labeling of neurons in P6 mice is possible is novel and demonstrates that cFos can be robustly expressed in the brains of neonatal mice. This transgenic system can therefore be an effective and advantageous tool for the investigation of early life neuronal activation within the brain and its consequences later in life.

An important consideration when pursuing activity-dependent labeling is the particular gene locus governing CRE expression. Immediate early genes (IEGs) represent a well-described connection between neuronal activity and subsequent gene expression changes (38), and thus provide a useful strategy for targeting active cell populations for genetic access. While several IEGs are expressed in the brain, cFos is known to be expressed in the neonatal rodent brain, and this expression in the PVT is dependent upon an ongoing stimulus (25,39). cFos has been shown to be directly involved in the long-term consequences of neuronal activation on transcriptional and circuit-level changes (40). Unlike other IEGs that act rapidly via direct influences on synapses and cellular function, cFos functions through more protracted pathways via regulation of downstream target genes (25, 39). Additionally, the role of cFos in memory is well-established, including a role in mediating responses following acquisition of contextual memories (39). While cFos is expressed following activation in both excitatory and inhibitory neurons and therefore cannot distinguish between the two, the vast majority of the PVT neurons are excitatory glutamatergic neurons (41,42). Therefore, the lack of distinction between excitatory and inhibitory neurons is not of consequence in this context. These characteristics highlight cFos as an excellent tool for understanding brain activity in early life.

Using TRAP2, we find that the PVT is prominently activated during exposure to ELA as compared to normal rearing in male mice. By contrast, in females, there is little additional apparent neuronal activation in ELA-versus control-reared groups. In both sexes, the selectivity of PVT engagement by early-life experiences is striking, as other brain regions related to stress and reward contain few experience-engaged neurons, and this activation does not differ between control and ELA mice.

In adult rodents, the PVT is activated by stimuli of both positive valence (e.g. drugs of abuse; 43) and negative valence (e.g. footshock; 44), but activation of the PVT by emotionally salient stimuli early in life is not well characterized. We have previously identified PVT activation in 9-day old rat pups by sensory input from their mothers, a positive emotionally salient experience (45). Here we show that the PVT is activated by ELA taking place during postnatal days 2-9 in mice. This is important, because adult studies demonstrate that activation of the PVT by stressful events influences responses to an additional stress later in life (45,46). Our demonstration that the PVT is selectively engaged by ELA renders this brain region a candidate for mediating the long-lasting consequences of ELA on future responses to stress throughout life.

In addition to altered stress responses (32,47,48), ELA leads to deficits in motivated reward behaviors (11,48). Activation of subsets of PVT neurons may mechanistically contribute to these consequences: The PVT is required for the retrieval of remote emotionally salient experiences (i.e. those that have occurred >24 hrs. ago) and their influence on the regulation of motivated behaviors in adult rodents (45,50,51). The PVT and its specific projections contribute to a variety of reward-related behaviors, including motivation for feeding (52), binge ethanol drinking (53) and heroin relapse (54). Disruptions in similar reward-related behaviors characterize numerous mood disorders, including substance use disorders and depression, for which ELA is a risk factor (2,5,55). Thus, the robust and differential engagement of the PVT during ELA reported here suggests that the PVT may encode these early-life experiences and utilize this information later in life to impact reward-related behaviors. These hypotheses will be subjects of future studies.

We find that ELA exerts the largest increase in activation in specific subregions of the PVT. The specific topology of this differential activation sheds light on the potential connectivity and functions of ELA-engaged PVT neurons: The posterior (p)PVT sends strong projections to regions including the ventromedial nucleus accumbens shell, central amygdala (CeA), basolateral amygdala (BLA) and bed nucleus of the stria terminalis, whereas the anterior (a)PVT projects in a more diffuse manner to regions including the dorsomedial nucleus accumbens shell, suprachiasmatic nucleus, and ventral subiculum (23,56,57). In addition to this heterogeneity in projection patterns, different contributions to motivated behaviors are attributed to the aPVT versus pPVT. For example, inactivation of the aPVT, but not pPVT, decreases sucrose seeking when an expected sucrose reward is omitted (58). Similarly, injection of neurotensin into the pPVT, but not aPVT, is sufficient to curb excessive ethanol consumption in rats (59). A role in regulation of responses to chronic or repeated stress has also been identified exclusively for the pPVT (18,60). Therefore, the distinct topological distribution of neuronal activation by ELA may suggest specific roles and projection targets that warrant further investigation.

Our finding that CRFR1-expressing PVT neurons are preferentially activated by ELA is intriguing. The receptor is well expressed during the first week of life in the rodent (61) and was abundantly expressed in both the male and female PVT in our findings. Why and how might ELA augment activation of CRFR1 neurons? As shown in other brain regions, ELA might increase CRH expression and release in the PVT, activating receptor-expressing neurons. CRH release regulates CRFR1 expression in a biphasic manner (62), such that it is conceivable that more cells in the PVT express CRFR1 above detection threshold in ELA versus control animals. The augmented activation of CRFR1-neurons by ELA is notable, because CRFR1 plays important roles in responses to stress, including mediating changes in reward-related behaviors following stress (63-66). Further analysis revealed a significant interaction between rearing condition and sex in the aPVT, whereby the effect of ELA on proportion of TRAPed aPVT neurons that express CRFR1 was significantly more pronounced in females as compared to males (Fig. S2). This, combined with the finding that overall PVT activation is greater in control-reared females as compared to control-reared males, suggests sex-dependent mechanisms of PVT activation in early life. Future studies will determine whether these CRFR1 neurons mediate the influence of ELA on adult reward behaviors.

In conclusion, the present studies describe a novel use of activity-dependent genetic tagging to demonstrate robust and selective activation of the PVT during ELA. Compared to typical rearing, PVT neurons activated during ELA are significantly more likely to express the receptor to CRH, CRFR1. Establishing the functional role of these ELA-engaged PVT neurons will be a crucial next step toward determining their role in disruptions of reward-related behaviors induced by ELA and will provide important information toward understanding the mechanisms underlying the consequences of ELA on mental health.

## Acknowledgements and Disclosures

This work was supported by National Institutes of Health grants F30 MH126615, T32 DA050558, T32 GM008620, and P50 MH096889. We thank Qinxin Ding and Manlin Shao for technical assistance. The authors report no biomedical financial interests or potential conflicts of interest.

## Supplementary info

**Supplementary Fig 1.**
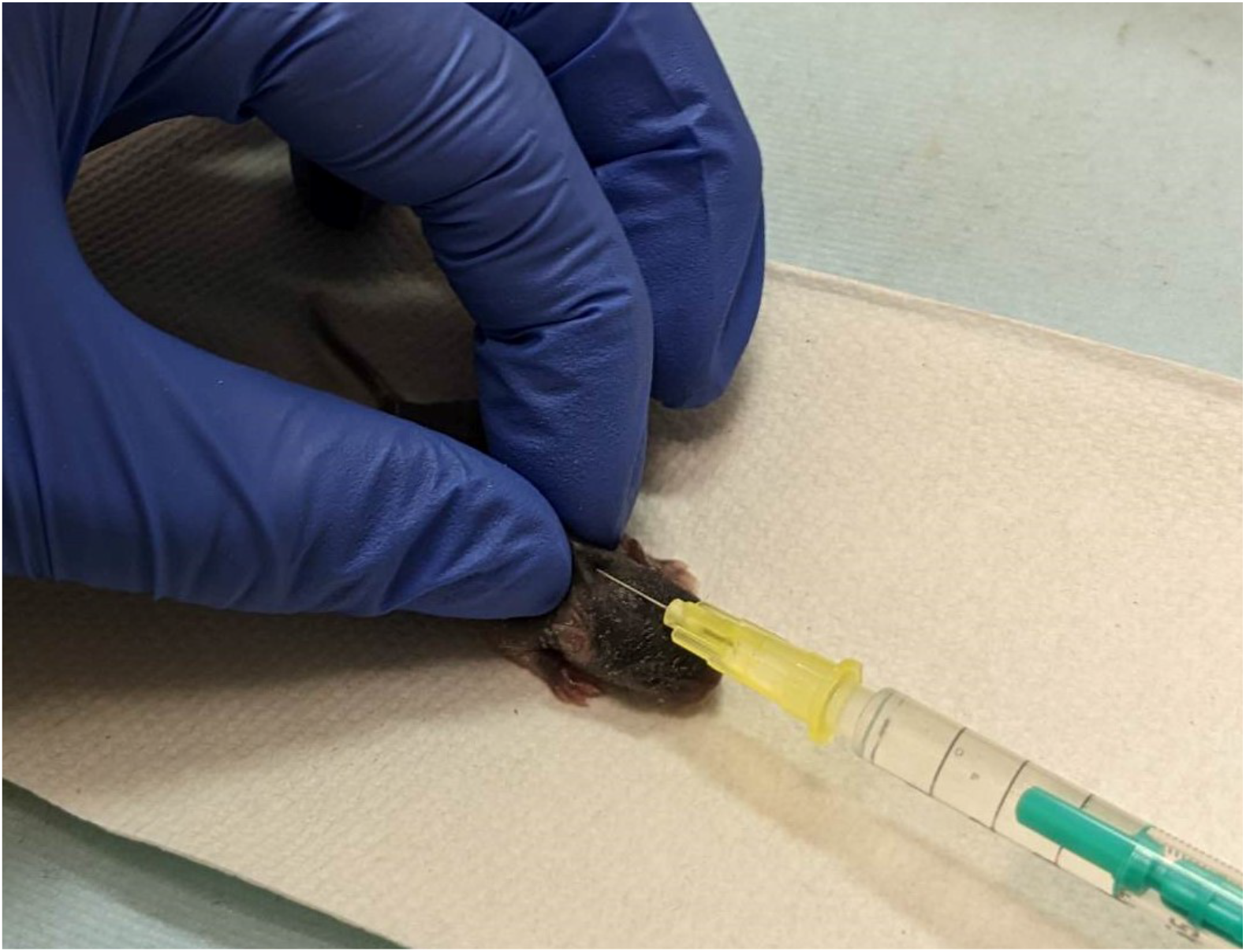
Subcutaneous injection of tamoxifen on P6. On the morning of P6, pups are briefly removed from their home cage (<3 min.), placed on a heating pad, and injected with 150 mg/kg tamoxifen dissolved in corn oil. The skin on the back of the neck is pinched to create a fold, into which a 30g needle is inserted for the injection. All live-animal procedures demonstrated here have been reviewed and approved by the Institutional Animal Care and Use Committee at UC Irvine.

**Supplementary Fig 2.**
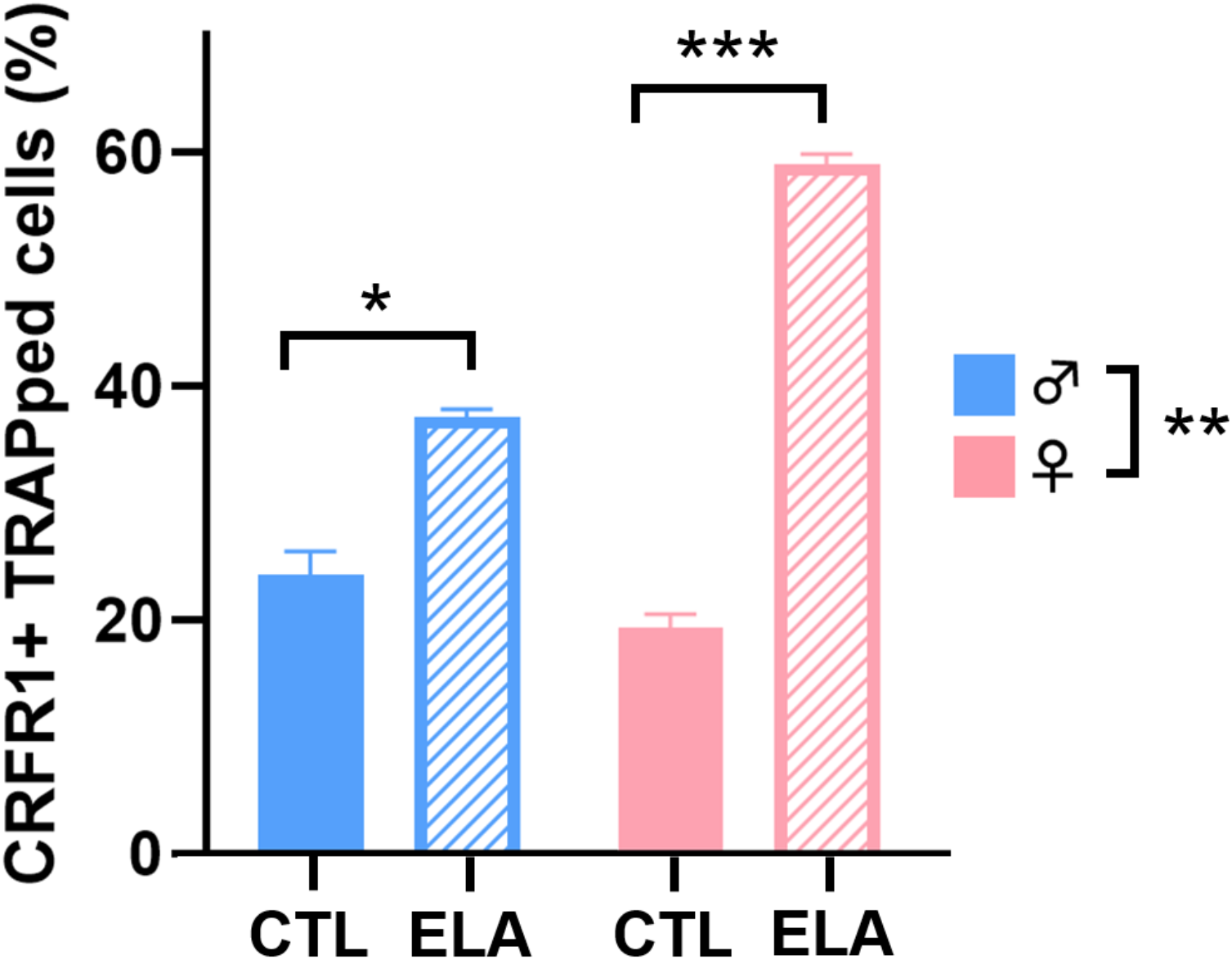
Proportion of aPVT neurons expressing CRFR1^+^ TRAPed neurons in CTL and ELA males and females. Comparing the proportion of TRAPed aPVT neurons that express CRFR1 revealed a significant interaction between rearing condition and sex (p = 0.0024; n = 5-6 mice/group; Two-way mixed model ANOVA with Sidak *post hoc* test). ***, p < 0.001; **, p < 0.01; *, p < 0.05. All values shown as mean ± SEM.

## References

1. American Psychiatric Association (2018): Stress in America Survey: Stress and Generation Z. Washington, DC: American Psychiatric Publishing.

2. Danese A (2017): Psychoneuroimmunology of early-life stress: the hidden wounds of childhood trauma? Neuropsychopharmacology 42:99–114.

3. Short A, Baram TZ (2019): Adverse early-life experiences and neurological disease: Age-old questions and novel answers. Nat Rev Neurol 15:657–669.

4. Silvers JA, Goff B, Gabard-Durnam LJ, Gee DG, Fareri DS, Caldera C, et al. (2017): Vigilance, the Amygdala, and Anxiety in Youths With a History of Institutional Care. Biol Psychiatry Cogn Neurosci Neuroimaging 2:493–501.

5. Green JG, Mclaughlin KA, Berglund PA, Gruber MJ, Sampson NA, Zaslavsky AM, et al. (2010): Childhood Adversities and Adult Psychiatric Disorders in the National Comorbidity Survey Replication I: Associations With First Onset of DSM-IV Disorders. Arch Gen Psychiatry 67:113–123.

6. Hackman DA, Farah MJ (2009): Socioeconomic status and the developing brain. Trends Cogn Sci 13:65–73.

7. McLaughlin KA, Weissman D, Bitrán D (2019): Childhood Adversity and Neural Development: A Systematic Review. Annu Rev Dev Psycho 1:277–312.

8. Callaghan BL, Sullivan RM, Howell B, Tottenham N (2014): The international society for developmental psychobiology Sackler symposium: Early adversity and the maturation of emotion circuits-A cross-species analysis. Dev Psychobiol 56:1635–1650.

9. Chen Y, Baram TZ (2016): Toward Understanding How Early-Life Stress Reprograms Cognitive and Emotional Brain Networks. Neuropsychopharmacology 41:197–206.

10. Bolton JL, Molet J, Regev L, Chen Y, Rismanchi N, Haddad E, et al. (2018): Anhedonia Following Early-Life Adversity Involves Aberrant Interaction of Reward and Anxiety Circuits and Is Reversed by Partial Silencing of Amygdala Corticotropin-Releasing Hormone Gene. Biol Psychiatry 83:137–147.

11. Bolton JL, Ruiz CM, Rismanchi N, Sanchez GA, Castillo E, Huang J, et al. (2018): Early-life adversity facilitates acquisition of cocaine self-administration and induces persistent anhedonia. Neurobiol Stress 8:57–67.

12. Levis SC, Bentzley BS, Molet J, Bolton JL, Perrone CR, Baram TZ, et al. (2020): On the early-life origins of vulnerability to opioid addiction. Mol Psychiatry 26:4409–4416.

13. Molet J, Heins K, Zhuo X, Mei YT, Regev L, Baram TZ, et al. (2016): Fragmentation and high entropy of neonatal experience predict adolescent emotional outcome. Transl Psychiatry 6:e702.

14. Oltean LE, Şoflău R, Miu A, Szentágotai-Tătar A (2022): Childhood adversity and impaired reward processing: A meta-analysis. Child Abuse Negl 105596.

15. DeNardo LA, Liu CD, Allen WE, Adams EL, Friedmann D, Fu L, et al. (2019): Temporal evolution of cortical ensembles promoting remote memory retrieval. Nat Neurosci 22:460–469.

16. Barson JR, Mack NR, Gao WJ (2020): The Paraventricular Nucleus of the Thalamus Is an Important Node in the Emotional Processing Network. Front Behav Neurosci 14:598469.

17. Hsu DT, Kirouac GJ, Zubieta JK, Bhatnagar S (2014): Contributions of the paraventricular thalamic nucleus in the regulation of stress, motivation, and mood. Front Behav Neurosci 8:1–10.

18. Bhatnagar S, Viau V, Chu A, Soriano L, Meijer OC, Dallman F (2000): A Cholecystokinin-Mediated Pathway to the Paraventricular Thalamus Is Recruited in Chronically Stressed Rats and Regulates Hypothalamic-Pituitary-Adrenal Function. J Neurosci 20:5564–5573.

19. Otis JM, Zhu M, Namboodiri VMK, Cook CA, Kosyk O, Matan AM, et al. (2019): Paraventricular thalamus projection neurons integrate cortical and hypothalamic signals for cue-reward processing. Neuron 103:423–431.

20. Choi EA, McNally GP (2017): Paraventricular thalamus balances danger and reward. J Neurosci 37:3018–3029.

21. Choi EA, Jean-Richard-dit-Bressel P, Clifford CWG, McNally GP (2019): Paraventricular thalamus controls behavior during motivational conflict. J Neurosci 39:4945–4958.

22. Kirouac GJ (2015): Placing the paraventricular nucleus of the thalamus within the brain circuits that control behavior. Neurosci Biobeh Rev 56:315–329.

23. Li S, Kirouac GJ (2008): Projections from the Paraventricular Nucleus of the Thalamus to the Forebrain, With Special Emphasis on the Extended Amygdala. J Comp Neurol 259:263–287.

24. Dong X, Li S, Kirouac GJ (2017): Collateralization of projections from the paraventricular nucleus of the thalamus to the nucleus accumbens, bed nucleus of the stria terminalis, and central nucleus of the amygdala. Brain Struct Funct 222:3927–3943.

25. Fenoglio KA, Chen Y, Baram TZ (2006): Neuroplasticity of the Hypothalamic-Pituitary-Adrenal Axis Early in Life Requires Recurrent Recruitment of Stress-Regulating Brain Regions. J Neurosci 26:2434–2442.

26. Molet J, Maras PM, Avishai-Eliner S, Baram TZ (2014): Naturalistic rodent models of chronic early-life stress. Dev Psychobiol 56:1675–1688.

27. Chen Y, Molet J, Gunn BG, Ressler K, Baram TZ (2015): Diversity of reporter expression patterns in transgenic mouse lines targeting corticotropin-releasing hormone-expressing neurons. Endocrinol 156:4769–4780.

28. Schindelin J, Arganda-Carreras I, Frise E, Kaynig V, Longair M, Pietzsch T, et al. (2012): Fiji: An open-source platform for biological-image analysis. Nat Methods 9:676–682.

29. Gilles EE, Schultz L, Baram TZ (1996): Abnormal corticosterone regulation in an immature rat model of continuous chronic stress. Pediatr Neurol 15:114–119.

30. Ivy AS, Brunson KL, Sandman C, Baram TZ (2008): Dysfunctional nurturing behavior in rat dams with limited access to nesting material: A clinically relevant model for early-life stress. Neuroscience 154:1132–1142.

31. Bolton JL, Short AK, Simeone K, Daglian J, Baram TZ (2019): Programming of Stress-Sensitive Neurons and Circuits by Early-Life Experiences. Front Behav Neurosci 13:1–9.

32. Short AK, Thai CW, Chen Y, Kamei N, Pham AL, Birnie MT, et al. (2021): Single-Cell Transcriptional Changes in Hypothalamic Corticotropin-Releasing Factor–Expressing Neurons After Early-Life Adversity Inform Enduring Alterations in Vulnerabilities to Stress. Biol Psychiatry Global Open Sci.

33. Dubé CM, Molet J, Singh-Taylor A, Ivy A, Maras PM, Baram TZ (2015): Hyper-excitability and epilepsy generated by chronic early-life stress. Neurobiol Stress 2:10–19.

34. Ivy AS, Rex CS, Chen Y, Dubé C, Maras PM, Grigoriadis DE, et al. (2010): Hippocampal dysfunction and cognitive impairments provoked by chronic early-life stress involve excessive activation of CRH receptors. J Neurosci 30:13005–13015.

35. Peng J, Long B, Yuan J, Peng X, Ni H, Li X, et al. (2017): A quantitative analysis of the distribution of CRH neurons in whole mouse brain. Front Neuroanat 11:63.

36. Itoga CA, Chen Y, Fateri C, Echeverry PA, Lai JM, Delgado J, et al. (2019): New viral-genetic mapping uncovers an enrichment of corticotropin-releasing hormone-expressing neuronal inputs to the nucleus accumbens from stress-related brain regions. J of Comp Neurol 527:2474–2487.

37. Eghbal-Ahmadi M, Hatalski CG, Lovenberg TW, Avishai-Eliner S, Chalmers DT, Baram TZ (1998): The developmental profile of the corticotropin releasing factor receptor (CRF2) in rat brain predicts distinct age-specific functions. Dev Brain Res 107:81–90.

38. Pinaud R, Tremere LA, De Weerd P (2005): Critical Calcium-Regulated Biochemical and Gene Expression Programs Involved in Experience-Dependent Plasticity. Plasticity in the Visual System: From Genes to Circuits. Boston: Springer, pp 153–180.

39. Gallo FT, Katche C, Morici JF, Medina JH, Weisstaub NV (2018): Immediate early genes, memory and psychiatric disorders: Focus on c-Fos, Egr1 and Arc. Front Behav Neurosci 12:1–16.

40. West AE, Griffith E, Greenberg ME (2002): Regulation of transcription factors by neuronal activity. Nat Rev Neurosci 3:921–931.

41. Frassoni C, Spreafico R, Bentivoglio M (1997): Glutamate, aspartate and co-localization with calbindin in the medial thalamus. An immunohistochemical study in the rat. Exp Brain Res 115:95–104.

42. Gupta A, Gargiulo AT, Curtis GR, Badve PS, Pandey S, Barson JR (2018): Pituitary adenylate cyclase-activating polypeptide-27 (PACAP-27) in the thalamic paraventricular nucleus is stimulated by ethanol drinking. Alcohol. Clin. Exp. Res 42:1650–1660.

43. Millan EZ, Ong ZY, McNally GP (2017): Paraventricular thalamus: Gateway to feeding, appetitive motivation, and drug addiction. Prog Brain Res 235:113–137.

44. Gao C, Leng Y, Ma J, Rooke V, Rodriguez-Gonzalez S, Ramakrishnan C, et al. (2020): Two genetically, anatomically and functionally distinct cell types segregate across anteroposterior axis of paraventricular thalamus. Nat Neurosci 23:217–228.

45. Bhatnagar S, Huber R, Nowak N, Trotter P (2002): Lesions of the posterior paraventricular thalamus block habituation of hypothalamic-pituitary-adrenal responses to repeated restraint. J Neuroendocrinol 14:403–410.

46. Do-Monte FH, Quinõnes-Laracuente K, Quirk GJ (2015): A temporal shift in the circuits mediating retrieval of fear memory. Nature 519:460–463.

47. Bolton JL, Short AK, Othy S, Kooiker CL, Shao M, Gunn BG, et al. (2022): Early stress-induced impaired microglial pruning of excitatory synapses on immature CRH-expressing neurons provokes aberrant adult stress responses. Cell Rep 38:110600.

48. Brunton PJ, Russell JA (2010): Prenatal social stress in the rat programmes neuroendocrine and behavioural responses to stress in the adult offspring: Sex-specific effects. J Neuroendocrinol 22:258–271.

49. Kangas BD, Short AK, Luc OT, Stern HS, Baram TZ, Pizzagalli DA (2022): A cross-species assay demonstrates that reward responsiveness is enduringly impacted by adverse, unpredictable early-life experiences. Neuropsychopharmacology 47:767–775.

50. Keyes PC, Adams EL, Chen Z, Bi L, Nachtrab G, Wang VJ, et al. (2020): Orchestrating Opiate-Associated Memories in Thalamic Circuits. Neuron 107:1113–1123.

51. Padilla-Coreano N, Do-Monte FH, Quirk GJ (2012): A time-dependent role of midline thalamic nuclei in the retrieval of fear memory. Neuropharmacol 62:457–463.

52. Ye Q, Nunez J, Zhang X (2022): Oxytocin Receptor-Expressing Neurons in the Paraventricular Thalamus Regulate Feeding Motivation through Excitatory Projections to the Nucleus Accumbens Core. J Neurosci 42:3949–3964.

53. Levine OB, Skelly MJ, Miller JD, Rivera-Irizarry JK, Rowson SA, DiBerto JF, et al. (2021): The paraventricular thalamus provides a polysynaptic brake on limbic CRF neurons to sex-dependently blunt binge alcohol drinking and avoidance behavior in mice. Nat Commun 12:5080.

54. Giannotti G, Gong S, Fayette N, Herson PS, Ford CP, Peters J, et al. (2021): Extinction blunts paraventricular thalamic contributions to heroin relapse. Cell Rep 36:109605.

55. Levis SC, Baram TZ, Mahler SV (2021): Neurodevelopmental origins of substance use disorders: Evidence from animal models of early-life adversity and addiction. Eur J Neurosci 55:2170–2195.

56. Moga MM, Weis RP, Moore RY (1995): Efferent projections of the paraventricular thalamic nucleus in the rat. J Comp Neurol 359:221–238.

57. Vertes RP, Hoover WB (2008): Projections of the paraventricular and paratenial nuclei of the dorsal midline thalamus in the rat. J Comp Neurol 508:212–237.

58. Do-Monte FH, Minier-Toribio A, Quiñones-Laracuente K, Medina-Colón EM, Quirk GJ (2017): Thalamic Regulation of Sucrose Seeking during Unexpected Reward Omission. Neuron 94:388–400.

59. Pandey S, Badve PS, Curtis GR, Leibowitz SF, Barson JR (2019): Neurotensin in the posterior thalamic paraventricular nucleus: inhibitor of pharmacologically relevant ethanol drinking. Addict Biol 24:3–16.

60. Heydendael W, Sharma K, Iyer V, Luz S, Piel D, Beck S, et al. (2011): Orexins/hypocretins act in the posterior paraventricular thalamic nucleus during repeated stress to regulate facilitation to novel stress. Endocrinol 152:4738–4752.

61. Avishai-Eliner S, Gilles EE, Eghbal-Ahmadi M, Bar-El Y, Baram TZ (2001): Altered regulation of gene and protein expression of hypothalamic-pituitary-adrenal axis components in an immature rat model of chronic stress. J Neuroendocrinol 13:799–807.

62. Brunson KL, Grigoriadis DE, Lorang MT, Baram TZ (2002): Corticotropin-releasing hormone (CRH) downregulates the function of its receptor (CRF1) and induces CRF1 expression in hippocampal and cortical regions of the immature rat brain. Exp Neurol 176:75–86.

63. Kreibich AS, Briand L, Cleck JN, Ecke L, Rice KC, Blendy JA (2009): Stress-induced potentiation of cocaine reward: a role for CRF R1 and CREB. Neuropsychopharmacology 34:2609–2617.

64. Chen NA, Jupp B, Sztainberg Y, Lebow M, Brown RM, Kim JH, et al. (2014): Knockdown of CRF1 receptors in the ventral tegmental area attenuates cue- and acute food deprivation stress-induced cocaine seeking in mice. J Neurosci 34:11560–11570.

65. Vranjkovic O, Van Newenhizen EC, Nordness ME, Blacktop JM, Urbanik LA, Mathy JC, et al. (2018): Enhanced CRFR1-dependent regulation of a ventral tegmental area to prelimbic cortex projection establishes susceptibility to stress-induced cocaine seeking. J Neurosci 38:10657–10671.

66. Lemos JC, Wanat MJ, Smith JS, Reyes BAS, Hollon NG, Van Bockstaele EJ, et al. (2012): Severe stress switches CRF action in the nucleus accumbens from appetitive to aversive. Nature 490:402–406.

